# The RNA modification N^6^-methyladenosine restrains SIRPα to license myeloid efferocytosis in atherosclerosis

**DOI:** 10.64898/2026.09.19.749544

**Authors:** Aneesh Kallapur, Xiang Li, Elena Rodriguez-Sanchez, Zhengyi Zhang, Aniketa Sinha, Lijing Cheng, Dan Wang, Jing Gu, Jason Kim, Xiaohui Wu, Wei Li, Ya Cui, Xin He, Chuan He, Tamer Sallam

**Affiliations:** Division of Cardiology, Department of Medicine, University of California, Los Angeles, Los Angeles, CA, USA; Department of Physiology, University of California, Los Angeles, Los Angeles, CA, USA; Molecular Biology Institute, University of California, Los Angeles, Los Angeles, CA, USA; Department of Human Genetics, The University of Chicago, Chicago, IL, USA; Division of Computational Biomedicine, Biological Chemistry, University of California, Irvine, Irvine, CA, USA; Department of Chemistry and Institute for Biophysical Dynamics, The University of Chicago, Chicago, IL, USA; Howard Hughes Medical Institute, The University of Chicago, Chicago, IL, USA; Department of Biochemistry and Molecular Biology, The University of Chicago, Chicago, IL, USA

## Abstract

**Background:** Macrophages participate in atherosclerotic plaque development, progression and resolution, integrating lipid and inflammatory cues to balance tissue damage and repair. Despite considerable understanding of macrophage function in atherosclerosis, the underlying regulatory mechanisms governing these responses continue to evolve. Emerging evidence implicates epitranscriptomic gene regulation by the RNA modification N^6^-methyladenosine (m^6^A) in control of context specific immune responses, but its role in atherosclerosis is not well-defined.

**Methods:** We integrated m^6^A quantitative trait loci (m^6^A QTLs) with Coronary Artery Disease (CAD) GWAS summary statistics. Using Oxford Nanopore direct RNA sequencing and MeRIP-seq, we constructed a high-resolution map of m^6^A modifications in macrophages. To determine the functional consequences of myeloid m^6^A loss *in-vivo*, we generated a myeloid-specific loss-of-function mouse model of m^6^A (*Mettl14*^f/f^ LysM^Cre^) and performed atherosclerosis studies using bone marrow transplantation into irradiated *Ldlr*^--/-^ mice. Atherosclerosis lesions were analyzed using light microscopy and immunofluorescence imaging, with complementary *in-vitro* mechanistic studies performed in murine bone marrow derived macrophages (BMDMs).

**Results:** m^6^A quantitative trait loci (m^6^A -QTLs) were enriched in CAD GWAS loci relevant to immune and lipid function, implicating m^6^A regulation in human CAD genetics. Using Oxford Nanopore direct RNA sequencing in conjunction with public MeRIP-sequencing data, we generated a single-base resolution, stoichiometric map of m^6^A modification in macrophages, nominating genes central to macrophage biology in atherosclerosis as high-confidence m^6^A targets. Myeloid specific deletion of the m^6^A “writer” complex subunit *Mettl14* drove the development of larger, more necrotic and unstable atherosclerotic plaques on an *Ldlr*^--/-^ background compared with *Mettl14*^f/f^ controls, establishing an atheroprotective role for myeloid m^6^A. Loss of myeloid *Mettl14* impaired macrophage efferocytosis in atherosclerotic plaques, in a dexamethasone-induced apoptosis in the thymus and *in-vitro*. Molecularly, m^6^A methylation of the “don’t eat me” antiphagocytic receptor SIRPα RNA suppressed its protein expression.

**Conclusions:** These findings identify the myeloid m^6^A - SIRPα axis as a novel homeostatic regulator of efferocytosis and establish epitranscriptomic remodeling as a crucial layer of macrophage regulation in atherosclerosis.

## Introduction

Immune cells, particularly macrophages, play multi-faceted roles in atherosclerosis development and progression. Acting as important conduits between environmental cues and gene regulatory mechanisms, macrophage take up modified low-density lipoprotein (LDL) cholesterol, triggering downstream inflammatory cascades that drive plaque progression^1^. Macrophages also play a critical atheroprotective roles through the phagocytic clearance of apoptotic cells, a process known as efferocytosis, to promote inflammation resolution and limit plaque necrosis^2^. The dynamic regulation of lipid metabolism, inflammatory signaling and pro-resolving responses by macrophages is therefore critical to the pathogenesis of atherosclerosis. While the control of these processes by lipid sensitive nuclear receptors such as Liver X Receptor (LXR)^3–6^ and the Peroxisome Proliferator-Activated Receptor (PPAR)^7,8^ family is established, the broader landscape of homeostatic regulatory mechanisms remains poorly defined.

Paque-rupture events, like myocardial infarction and stroke, represent some of the most devastating forms of cardiovascular disease. Human autopsy studies^9^ and prospective imaging trials^10^ have consistently shown that thin-cap fibroatheromas (TCFA), vulnerable plaques defined by a large necrotic core beneath a thin fibrous cap, are highly prone to rupture even independent of the underlying lesion burden. This vulnerability is thought to arise in part from defective macrophage efferocytosis within the plaque, a key driver of necrotic core expansion and lesion instability^11–15^. Human genetic evidence further indicates that defective efferocytosis causally contributes to CAD risk^2^, which has driven endeavors to target this process therapeutically^14,16,17^. In particular, ongoing efforts to target the anti-phagocytic “don’t eat me” CD47- SIRPα axis are in early-stage clinical trials^14,17^

Emerging evidence has implicated “epitranscriptomic” regulation by RNA modifications such as N^6^-methyladenosine (m^6^A) as context dependent modulators of gene expression, acting through control of mRNA stability, translation efficiency as well as modulation of chromatin accessibility and transcription^18^.These m^6^A marks, installed by the methyltransferase “writer” proteins including METTL3, METTL14 and WTAP, and removed by demethylase “eraser” proteins including FTO and ALKBH5, have been shown to be therapeutically tractable^19^ and are under investigation in clinical trials in acute leukemia. Beyond its integral roles in cellular development and stress responses^18^, m^6^A has emerged as a key regulator of innate immune function, with demonstrated roles in host-pathogen defense^20^, macrophage polarization^21^ and leukocyte differentiation^22^. Multiple studies have also implicated m^6^A as a brake on visceral lipid accumulation by fine-tuning lipid metabolism^23,24^. Although prior work has studied the role of m^6^A in atherosclerosis in the context of vascular cell types^25,26^, the canonical roles of m^6^A in host defense responses and leukocyte biology hint at a regulatory role of myeloid m^6^A in atherosclerosis. Furthermore, few studies have consolidated the role of m^6^A in the context of large-scale human genetic data complemented by murine mechanistic studies. Collectively, these observations raise the possibility that epitranscriptomic regulation of macrophage function may be involved in the progression of atherosclerosis.

A landmark technical development in the m^6^A field was MeRIP seq^27^, an antibody-based immunoprecipitation of m^6^A on mRNA followed by sequencing, to map m^6^A sites genome-wide. Despite its transformative impact, the method has several limitations that constrain both mechanistic understanding of m^6^A dynamics and precision targeting of individual modifications. MeRIP seq captures RNA fragments rather than the modified nucleotide itself, thus providing a 100-200 nucleotide window for the location of the putative modified base, making it challenging to design targeted base editing strategies^28^. Additionally, lack of stoichiometric information makes it difficult to determine which m^6^A methylated sites are methylated highly enough to influence transcript fate and thus biological effects. Finally, the anti-m^6^A antibody used in the immunoprecipitation cross-reacts with related RNA modifications, including m^6^A^m^, identifying false positive sites^29^. Methods that detect m^6^A at a single-base resolution with stoichiometric information, including Oxford Nanopore direct RNA sequencing^30^, and m^6^A-GLORI^31^ (glyoxal and nitrite-mediated deamination of unmethylated adenosines), address these limitations, but have only begun to be applied in physiologically relevant, disease-associated cell types^32^.

Here, we demonstrate that m^6^A is genetically associated with CAD risk in human GWAS and investigate its functional role in macrophage biology in atherosclerosis. Using Oxford Nanopore direct RNA sequencing, we map m^6^A modifications in macrophages in a single-base resolution. We show that myeloid m^6^A exerts atheroprotective effects *in-vivo*, promoting macrophage efferocytosis to limit plaque burden. Mechanistically, we demonstrate that m^6^A acts as a brake on anti-phagocytic responses. Collectively, these data demonstrate a homeostatic role of myeloid epitranscriptomic regulation in atherosclerosis.

## Results

### Human genetic evidence nominates a role for m^6^A in regulating immune cell responses in atherosclerosis

Molecular Quantitative Trait loci (mol-QTLs) including expression QTLs (eQTLs) and splice QTLs (sQTLs) have enabled systematic prioritization of candidate genes at non-coding GWAS loci. Similarly, recent efforts have mapped m^6^A -QTLs using MeRIP-seq in lymphoblastoid cells of 60 well-phenotyped Yoruba (YRI) individuals, demonstrating that m^6^A -QTLs were enriched in GWAS loci of multiple inflammatory traits, at frequencies comparable to other mol-QTLs.^33^. To determine whether specific CAD loci harbor m^6^A -QTL signals, we integrated m^6^A -QTLs with summary statistics from the largest published CAD GWAS meta-analysis to date^34^ (Figure 1A).

**Figure 1:**
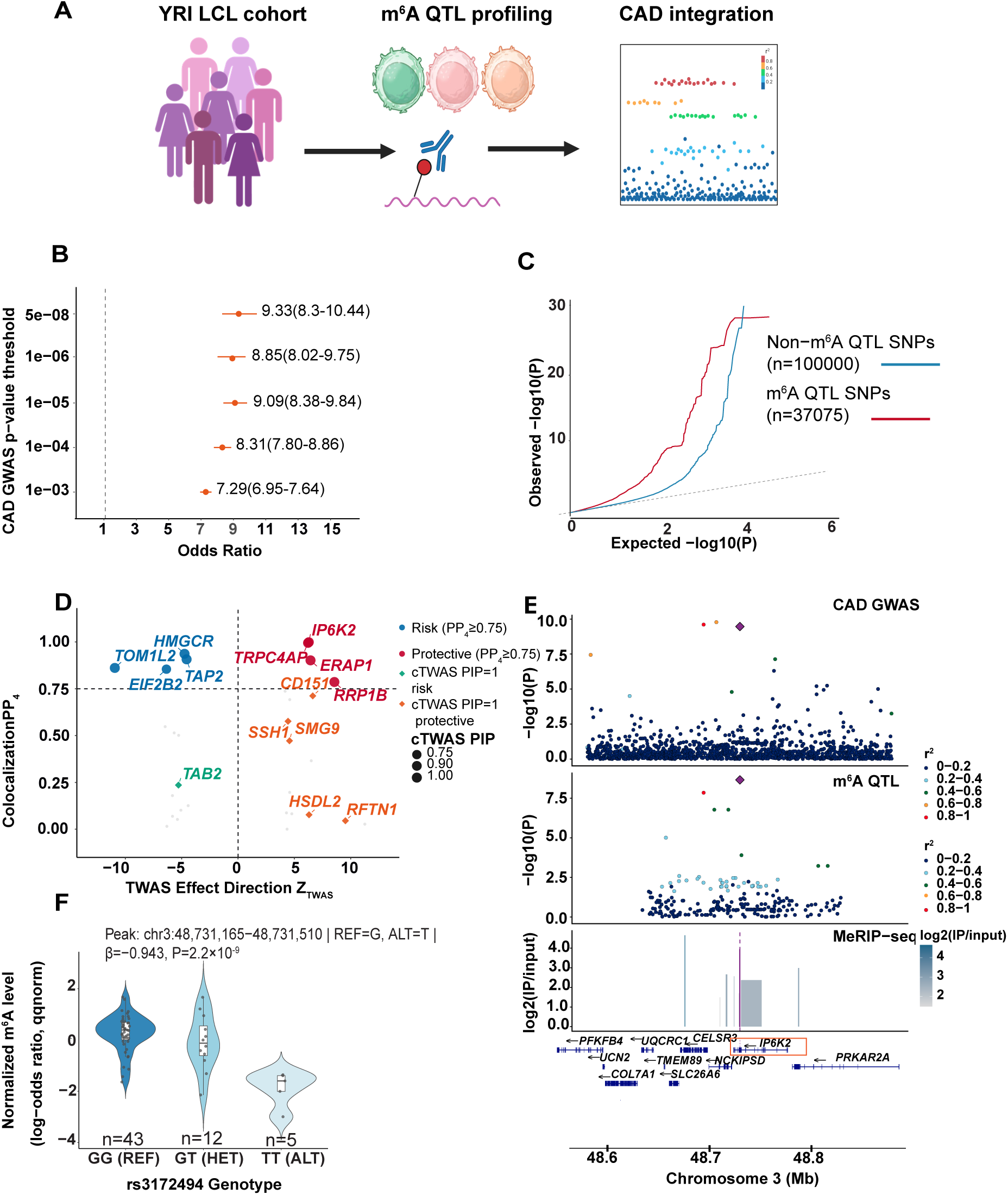
m^6^A-QTLs are enriched in CAD GWAS loci. (A) Schematic of MeRIP-seq in LCLs of 60 YRI individuals (schematic made with Biorender). (B) Forest plot showing Odds Ratio of enrichment of m^6^A-QTLs in CAD GWAS loci at various statistical thresholds. (C) Q-Q plot of enrichment of m^6^A-QTL SNPs relative to non m^6^A-QTL SNPs in CAD GWAS loci. (D)Plot of TWAS Z score vs Posterior probability of colocalization for m^6^A-QTLs.(E) Colocalization results of m^6^A-QTL and CAD GWAS SNPs. (F) Violin plot of the relative m^6^A with various rs3172494 genotypes Center line= median; box= interquartile range (IQR); whiskers=1.5xIQR. (G) LocusZoom plot of the *IP6K2* locus for CAD. Fisher’s exact test was used for statistical analysis in panel (c). YRI= Yoruba in Ibadan, Nigeria; LCL= Lymphoblastoid cell lines; QTL= Quantitative trait loci; CAD= Coronary Artery Disease; CI=Confidence interval, TWAS= Transcriptome wide association study; PIP= Posterior inclusion probability.

To assess enrichment of m^6^A QTLs in CAD GWAS loci, we annotated all SNPs in the CAD GWAS by their m^6^A QTL status, defining m^6^A QTLs as variants reaching FDR < 10%. This enrichment was first quantified by Fisher’s exact test comparing the proportion of m^6^A QTL SNPs among CAD genome-wide significant variants (p < 5×10⁻⁸) versus non-significant variants: 309 of 18,348 CAD- significant SNPs (1.7%) were m^6^A QTLs, compared to 36,766 of 20,054,722 non-significant SNPs (0.2%), yielding an odds ratio of 9.33 (95% CI: 8.30–10.44, p = 8.56×10⁻¹⁸¹). To assess whether this enrichment was driven by the most robustly associated CAD variants or reflected a more general phenomenon, we computed enrichment odds ratios across five GWAS significance thresholds ranging from p < 1×10⁻³ to p < 5×10⁻⁸ (Figure 1B). The odds ratio increased monotonically from 7.29 (95% CI: 6.95–7.64) at p < 1×10⁻³ to 9.33 (95% CI: 8.30–10.44) at the genome-wide significance threshold, with the proportion of SNPs that were m^6^A QTLs increasing from 1.3% to 1.7% across the same range. This dose-response relationship reflects genuine biological enrichment as opposed to linkage disequilibrium contamination or statistical artefact. To visualize this enrichment across the full allelic spectrum, we generated quantile-quantile (Q-Q) plots stratified by m^6^A QTL status (Figure 1C).m^6^A-QTL SNPs deviated more strongly from the null diagonal than non-m^6^A QTL SNPs, indicating systematic enrichment of m^6^A-QTLs in CAD GWAS loci.

To identify specific genes whose altered m^6^A modification mediates CAD susceptibility, we integrated pre-trained genetic prediction models of m^6^A abundance^33^ with CAD GWAS summary statistics in a transcriptome wide association study (m^6^A-TWAS) framework. Across 918 cis- heritable m^6^A peaks, we identified 34 transcriptome-wide significant loci passing strict Bonferroni correction (P_TWAS_<5.45x10^-5^). Subsequent Bayesian colocalization analysis confirmed that 8 of these associations are driven by shared causal variants (PP_H4_≥0.75) rather than incidental linkage disequilibrium. These high-confidence loci span distinct biological pathways including cholesterol biosynthesis (*HMGCR*), antigen presentation (*TAP2, ERAP1*), endosomal trafficking (*TOM1L2*) as well as inositol signaling and apoptosis (*IP6K2)*.

A key limitation of sequential TWAS and colocalization is that both methods test genes independently, making them susceptible to false positives arising from linkage disequilibrium between nearby genes. To account for this, we utilized “causal TWAS” (cTWAS)^35^, a Bayesian fine-mapping framework that jointly models all genes and SNPs within an independent LD region simultaneously, assigning posterior inclusion probabilities (PIPs), the Bayesian posterior probability that molecular traits affect the phenotype of interest.

cTWAS confirmed all eight colocalized loci with PIPs ranging from 0.76-1.00. Beyond the eight m^6^A sites in the colocalized loci, cTWAS identified 15 additional m^6^A sites with PIP≥0.8 which were missed by colocalization analysis, of which six had PIP=1.00. Of these, *TAB2* is a key adapter protein in the NF-κB innate immune signaling cascade^36^, and *SSH1* encodes a phosphatase regulating cytoskeletal dynamics in macrophages^37^. To jointly represent the strength of colocalization, fine-mapping confidence and direction of effect for each locus, we plotted TWAS z-scores (reflecting whether higher m^6^A increases or decreases CAD risk) against colocalization posterior probability (PP4) with point size proportional to cTWAS PIP (Figure 1D). High-confidence loci clustered at both positive and negative TWAS z-scores, indicating that m^6^A modification is risk-promoting at some loci (*IP6K2, TRPC4AP, ERAP1, RRP1B*) and protective at others (*HMGCR, TAP2, TOM1L2, EIF2B2*).

The strongest colocalization signal emerged at the *IP6K2* locus (cTWAS PIP 0.968, PP_H4_=0.999, Z_TWAS_=+6.28, P_TWAS_-3.15x10^-7^), where a single variant rs3172494 drives both the m^6^A-QTL and CAD GWAS associations with high confidence (per-SNP PP_4_=0.909).The rs3172494 alternate allele (T) was associated with significantly reduced m^6^A abundance on *IP6K2* transcripts (β_QTL_=- 0.943, p_QTL_=2.2x10^-9^, FDR=8.7x10^-6^) and a concordant reduction in CAD risk (β_GWAS_=-0.051, p_GWAS_=3.44x10^-10^), demonstrating that genetically lower m^6^A methylation at this locus is protective against CAD (Fig 1E). *IP6K2* encodes Inositol Hexakisphosphate Kinase 2, which synthesizes the Inositol pyrophosphate metabolite 5-IP7, a pathway required for p53 mediated apoptosis^38,39^ and implicated in NF-κB and IFN-β activation^38,40^

Collectively, these data provide human genetic evidence from multiple analytical frameworks that m^6^A modifications are associated with CAD risk, at least in part through immune and lipid mediated mechanisms.

### A map of m^6^A modifications in macrophages using Oxford Nanopore Sequencing

To study the functional role of m^6^A in macrophages, we generated a myeloid-specific loss- of-function model of m^6^A by deleting *Mettl14*, an essential structural component of the methyltransferase “writer” complex, in myeloid cells (*Mettl14*^f/f^LysM^Cre^). Deletion of METTL14 was confirmed by western blot (Fig S1A) and RT-qPCR of the deleted exons in BMDMs (Bone marrow derived macrophages) isolated from *Mettl14^f/^*^f^ (hereafter referred to as WT) or *Mettl14*^f/f^ LysM^Cre^ (hereafter referred to as M-KO) mice (Fig S1B).

To generate a comprehensive, quantitative map of m^6^A modifications in macrophages, we performed Oxford Nanopore direct RNA sequencing on WT and M-KO BMDMs. This technology sequences native mRNA as it passes through a pore and infers modifications from characteristic shifts in electrical signal, yielding per-site stoichiometry at single nucleotide resolution (Fig 2A).

**Figure 2:**
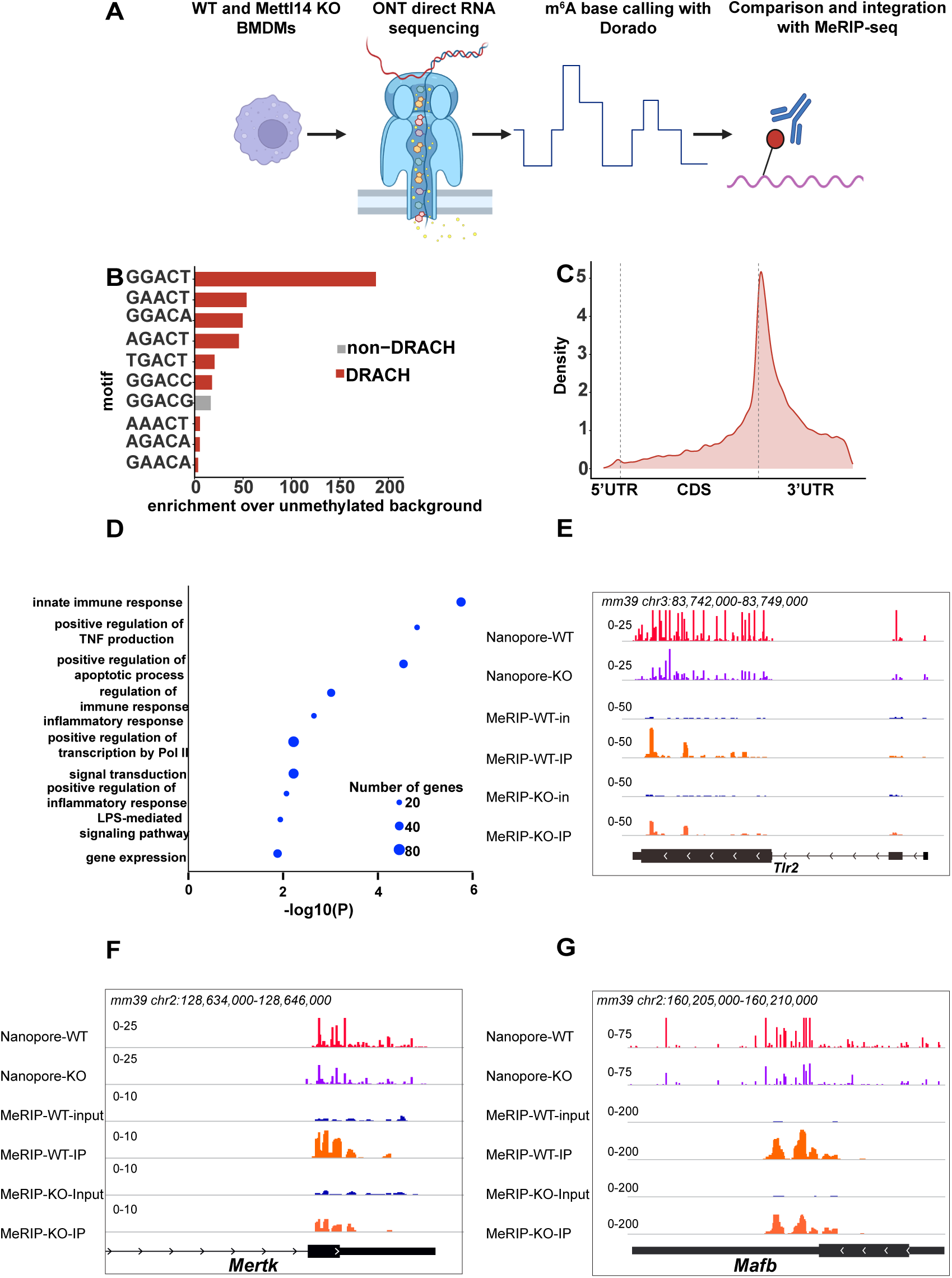
m^6^A target genes in macrophages are enriched for atherosclerosis relevant processes. (A) Schematic of Oxford Nanopore direct RNA sequencing in BMDMs (made with Biorender). (B) 5-mer motif enrichment over unmethylated background at putative Nanopore detected m^6^A sites. (C) Peak distribution plot of Nanopore detected m^6^A sites. (D) Gene ontology analysis of high-confidence m^6^A targets obtained by integrating Nanopore and MeRIP data.(E) Distribution of MeRIP and Nanopore peaks in the *Tlr2* gene. (F) Distribution of MeRIP and Nanopore peaks in the *Mertk* gene (G) Distribution of MeRIP and Nanopore peaks in the *Mafb* gene.

Unbiased 5-mer motif analysis identified the canonical m^6^A DRACH motifs (D = A/G/U, R = A/G, H = A/C/U) as the most enriched sequences genome-wide, with the most prominently enriched motif relative to unmethylated background being GGACT (Fig 2B, S1C). Genome-wide m^6^A distribution was concentrated at the 3’UTR, and peaked sharply at the stop-codon-3’UTR junction, the canonical topology of m^6^A^41^ (Fig 2C,S1D).

We defined bona-fide *Mettl14* dependent m^6^A sites as either (i) those with WT stoichiometry ≥ 50% and ≥20 percentage point reduction in M-KO or (ii) those with statistically significant reduction in M-KO (Fisher’s exact test FDR <0.05) with ≥50% relative or ≥5% absolute loss in M-KO. These criteria yielded 39,750 sites in 7,283 genes. To cross-validate and integrate these base calls, we reanalyzed a published MeRIP seq dataset performed in WT and *Mettl14* KO BMDMs^21^. We defined high-confidence m^6^A sites as bona-fide Nanopore sites falling within a MeRIP peak significantly lost in *Mettl14* KO BMDMs (FDR<0.05), which identified 3,567 dual- supported sites across 790 genes. Gene ontology analysis of this high-confidence set was strongly enriched for processes central to atherosclerosis, including inflammatory response, cytokine production, response to lipid and endocytosis (Fig 2D). Consistent with this, we identified high-stoichiometry *Mettl14*-dependent m^6^A sites in genes spanning multiple arms of atherosclerosis biology including inflammation (*Tlr2)*, efferocytosis (*Mertk*) and cell survival (*Mafb*)^42^ (Fig 2E,F,G).

Finally, we used our integrated data to compare detection between platforms. Across WT baseline stoichiometry thresholds, Nanopore recovered 40-71% of differentially methylated MeRIP peaks, with recovery rising as the threshold was relaxed (Fig S1E). Conversely, of bona-fide Nanopore sites, ∼63% were found outside any MeRIP peak, reflecting the greater sensitivity of per-molecule detection over antibody enrichment (Fig S1F).

Collectively, these data provide a quantitative, single-nucleotide map of the macrophage m^6^A methylome, demonstrate that *Mettl14*- dependent m^6^A marks genes across the breadth of atherosclerosis biology, and identify sites not captured by MeRIP-seq.

### Loss of myeloid *Mettl14* accelerates atherosclerosis and promotes plaque vulnerability

To investigate the contribution of myeloid m^6^A in atherosclerosis, we transplanted bone marrow from either WT or M-KO donors into lethally irradiated *Ldlr*^-/-^ recipient mice (Fig 3A). Successful engraftment was confirmed by PCR amplification of the Cre allele from peripheral blood DNA 8 weeks post-transplant (Fig S2A). Recipient mice were then placed on an atherogenic Western diet for 18 weeks to drive plaque formation.

**Figure 3:**
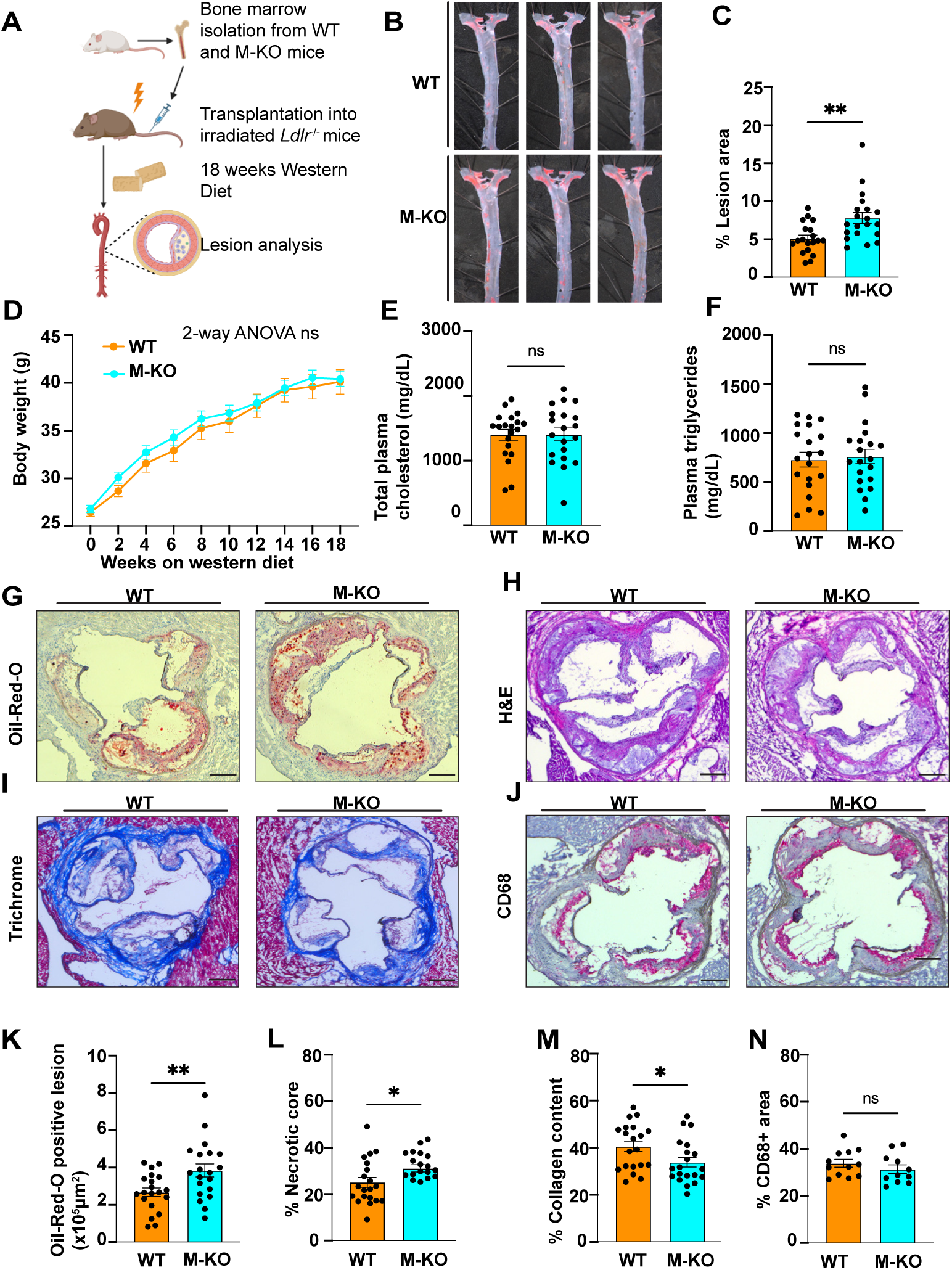
Loss of myeloid *Mettl14* leads to increased atherosclerosis. (A) Schematic of bone marrow transplantation (BMT) experiment. (B) Representative en-face staining of aorta of BMT mice. (C) Quantification of en-face lesion in the BMT study (n=19 in WT, n=20 in M-KO, one sample in WT excluded as an outlier after performing Grubb’s test, P<0.05). (D) Body weight gain in the BMT mice over the course of western diet feeding(n=20 per group).(E) Total plasma cholesterol measurement in the BMT mice(n=20/group). (F) Plasma triglyercide measurement in the BMT mice (n=20/group) (G) Representative Oil-Red-O images of aortic roots. (H) Representative Hematoxylin and Eosin (H&E) staining images of aortic roots. (I) Representative images of Masson’s Trichome staining of aortic roots. (J) Representative images of CD68 immunohistochemistry staining of aortic roots. (K) Quantification of Oil-Red-O staining of aortic roots (n=20/group). (L) Quantification of necrotic core percentage of aortic roots (n=20/group). (M) Quantification of percentage of collagen content of aortic roots (n=20/group). (N) Quantification of percentage of CD68 positive area in aortic roots (n=11/group). All data are presented as mean ± SEM. P values for (c), (e), (f), (k), (l), (m), and (n) were calculated using unpaired student’s t-test and for (d) with 2-way ANOVA *P<0.05, **P<0.01

M-KO bone marrow recipients developed significantly greater atherosclerotic burden than WT controls, as quantified by *en-*face Sudan Black B staining of whole aortae (Fig 3B,3C) and Oil Red O staining of aortic root cross-sections (Fig 3G,K). Importantly, this increase in plaque burden occurred in the absence of differences in body weight (Fig 3D), plasma cholesterol (Fig 3E) or plasma triglycerides (Fig 3F), demonstrating that the effect is independent of systemic metabolic perturbation and therefore intrinsic to the myeloid compartment. M-KO recipients also developed splenomegaly, with significantly larger spleen weights (Fig S2B,C) compared with WT controls, consistent with enhanced myeloid activation.

To define the morphological basis of increased plaque burden, we performed systematic histological characterization of aortic root sections. Hematoxylin and Eosin (H&E) staining revealed that M-KO recipients had a larger necrotic core to plaque ratio (Fig 3H,3L). Staining for lesional collagen with Masson Trichrome stain revealed that M-KO bone marrow transplanted mice had a significantly lower percentage of collagen within their plaques (Fig 3I,M), suggesting that loss of myeloid m^6^A promotes a vulnerable plaque phenotype. By contrast, lesional macrophage content, was equivalent between the genotypes (Fig 3J,N), as was macrophage proliferation, measured by the proportion of Ki67+ CD68+ nuclei (Fig S2E,F), indicating that the increased lesion burden was not explained by differences in macrophage accumulation or local proliferation.

We performed *in-vitro* assays in cultured WT and M-KO macrophages to determine whether the observed effects on plaque morphology could be attributed to intrinsic defects in macrophage cholesterol metabolism. M-KO peritoneal macrophages showed no difference from controls in foam cell formation after oxidized LDL treatment (Fig S3A) or cholesterol ester accumulation (Fig S3B) following acetylated LDL treatment, suggesting that altered macrophage cholesterol metabolism is unlikely to drive plaque progression in M-KO mice. We observed no evidence of systemic inflammation in *Mettl14* KO transplanted mice as assessed by the levels of pro-inflammatory cytokines that drive atherosclerosis including TNFα (Fig S3C), IL-18(Fig S3D), IL- 6(Fig S3E), RANTES (Fig S3F) and MCP-1 (Fig S3G).

Taken together, these findings establish that myeloid *Mettl14* plays an intrinsic atheroprotective role, limiting plaque growth and maintaining plaque stability.

### Loss of myeloid *Mettl14* impairs macrophage efferocytosis

Expansion of the necrotic core is canonically driven by defective efferocytosis, the phagocytic clearance of apoptotic cells within the plaque. We therefore performed Immunofluorescence (IF) staining for apoptotic cells using Terminal deoxynucleotidyl transferase dUTP Nick-End Labeling (TUNEL) alongside CD68 to mark macrophages. Quantification of TUNEL+ nuclei demonstrated an accumulation of apoptotic cells within M-KO plaques (Fig 4A,B). Accordingly, immunoblotting of splenic lysates showed an increase in cleaved caspase-3 protein in M-KO relative to WT (Fig 4D), further suggesting that loss of *Mettl14* lead to an accumulation of apoptotic cells in an atherosclerotic background.

**Figure 4:**
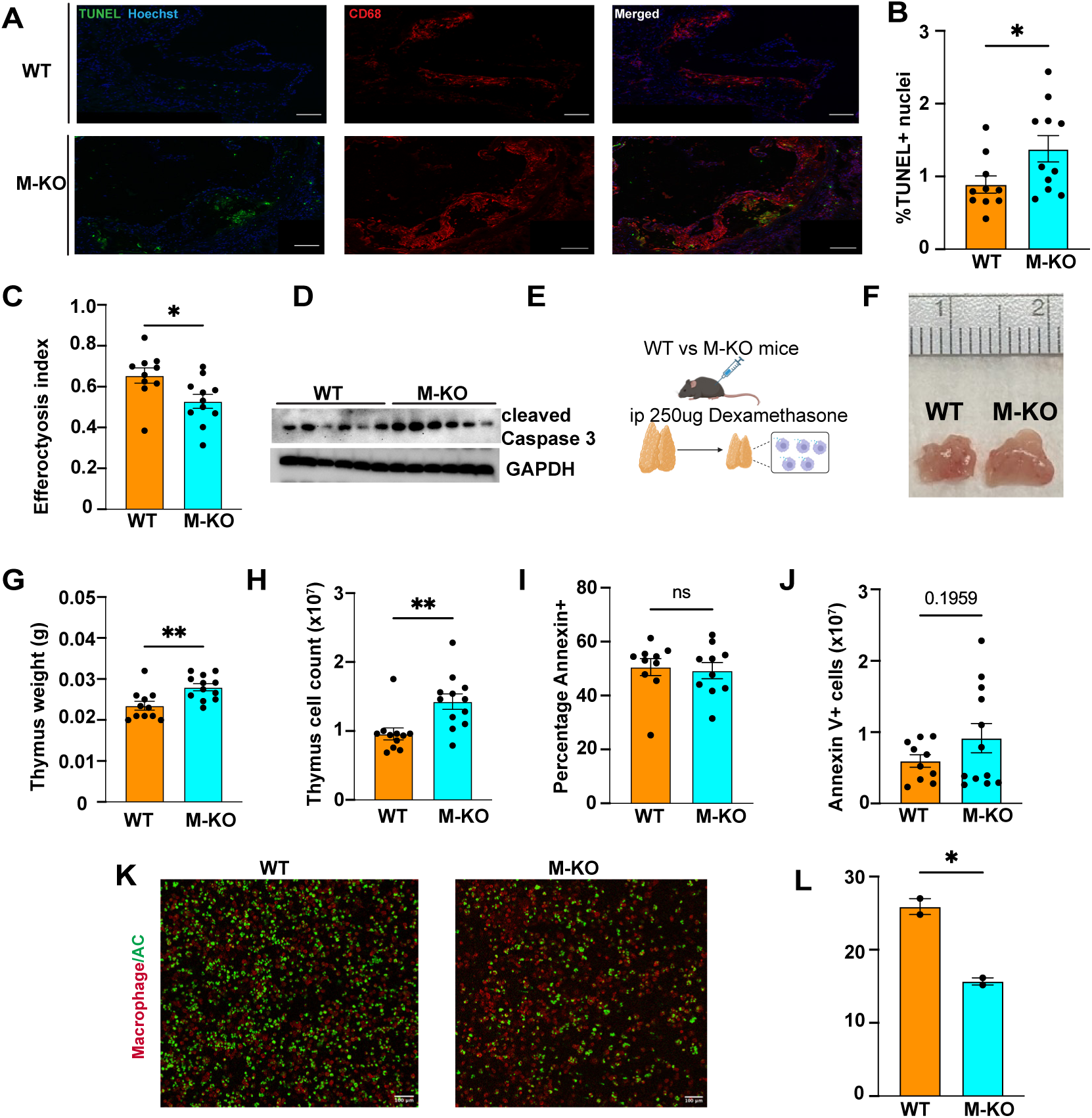
Loss of myeloid *Mettl14* leads to impaired macrophage efferocytosis. (A) Immunofluorescence images of aortic roots stained for TUNEL (green), CD68 (red) and Hoechst, scale bar=100μm. (B)Percentage of TUNEL positive nuclei in the plaque sections at the level of the aortic root (n=10 WT, 11 KO, 1 sample from WT formally excluded as an outlier using Grubb’s test). (C)Quantification of the efferocytosis index in plaque sections in the aortic root (n=10 WT, 11 KO) (D) Immunoblot for cleaved caspase 3 in the spleen of BMT mice. (E) Schematic of dexamethasone-thymus experiment. (F) Representative image of thymi from WT and M-KO mice after dexamethasone injection. (G) Weight of the thymi in WT and M-KO mice after dexamethasone injection (n=11 WT, 12 KO). (H) Cell count in WT and M-KO thymic lysates after dexamethasone injection (n=11 WT, 12 M-KO). (I) Percentage of Annexin V positive cells in WT and M-KO thymi after dexamethasone injection (n=10/group). (J) Number of Annexin V positive cells in WT and M-KO thymic lysates after dexamethasone injection (n=10 WT, 11 M-KO, one WT sample formally excluded as an outlier after Grubb’s test) . (K) Representative image of *in-vitro* efferocytosis (n=2 mice per group). scale bar=100μm, Macrophages are stained with wheat germ agglutinin conjugated to Texas Red and apoptotic cells are stained with Green CMFDA. (L) Quantification of *in vitro* efferocytosis, Data are presented as mean ± SEM. P values in (B), (C), (G), (H), (I), (J) and (L) were determined by

To quantify *in-situ* efferocytosis within the plaque, we utilized the validated “efferocytosis index”, the ratio of apoptotic cells (TUNEL+ nuclei) that are within the vicinity of macrophages (CD68+ area), which represent cells undergoing efferocytosis, to the apoptotic cells that are not within the vicinity of macrophages, representing unengulfed cells^43,44^. M-KO bone marrow recipients showed a significantly reduced efferocytosis index compared with controls (Fig 4C), establishing that myeloid *Mettl14* loss impairs efferocytosis within the atherosclerotic plaque.

To examine this finding in an orthogonal *in-vivo* context, we employed the dexamethasone-thymus model of efferocytosis. The corticosteroid dexamethasone is a potent inducer of thymic parenchymal cell apoptosis and triggers rapid, robust macrophage-mediated clearance, resulting in thymic involution^44,45^. We injected WT and M-KO mice with 250 μg of dexamethasone intraperitoneally and harvested the thymi 18 hours later(Fig 4E). M-KO mice exhibited significantly heavier (Fig 4F,G) and more cellular thymi (Fig 4H) compared with WT controls after Dexamethasone injection. Flow cytometric quantification (gating strategy Fig S4A) of Annexin V+ apoptotic cells revealed no significant difference in the proportion of apoptotic cells between genotypes (Fig 4I), but M-KO mice trended to harbor a greater absolute number of apoptotic cells (Fig 4J). Importantly, PBS injected WT and M-KO mice were indistinguishable at baseline in thymus weight (Fig S4B), cellularity (Fig S4C) and apoptotic cell proportion, confirming that the efferocytosis defect is dexamethasone dependent and not a consequence of altered thymic homeostasis.

Finally, we confirmed our findings with an *in-vitro* assay. WT and M-KO bone marrow derived macrophages (BMDMs) were co-incubated with CMFDA-labeled apoptotic Jurkat cells, and macrophages were counterstained with wheat germ agglutinin conjugated to a red fluorophore to delineate cell membranes. M-KO macrophages showed a significantly lower proportion of CMFDA+ apoptotic cell engulfment compared with WT controls, demonstrating a cell-intrinsic efferocytosis defect (Fig 4K,L).

Collectively, convergent evidence from atherosclerotic plaques, an *in-vivo* efferocytosis model and an *in-vitro* assay establish that myeloid *Mettl14* is cell-intrinsically required for efficient macrophage efferocytosis.

### SIRPα is directly regulated by m^6^A

To prioritize candidate driver m^6^A modified genes relevant to efferocytosis, we ranked all genes in the gene ontology pathways “efferocytosis”, “positive regulation of phagocytosis” and “negative regulation of phagocytosis” with site-level support from both Nanopore and MeRIP-seq data across six different evidence axes- three quantifying Nanopore methylation (number of m^6^A sites, WT stoichiometry and *Mettl14* dependent loss in M-KO) and three quantifying MeRIP support (number of peak, fold enrichment and differential log_2_ fold change) and combined them into an equal-weight composite score (Fig 5A). This analysis revealed that multiple genes in the efferocytosis and phagocytosis program exhibited high basal and differential m^6^A methylation, including canonical efferocytosis receptors such as *Mertk* and *Sirpa* as well as the Fcγ receptors *Fcgr1* and *Fcgr2b*. *Mertk* emerged as the lead candidate, exhibiting a relatively high number and stoichiometry of sites as well as differential methylation. *Mertk* mRNA (Fig S5A) and protein (Fig S5B) were significantly increased in *Mettl14* -KO mice, which suggests that defective efferocytosis in this context is likely being driven by other m^6^A target genes.

**Figure 5:**
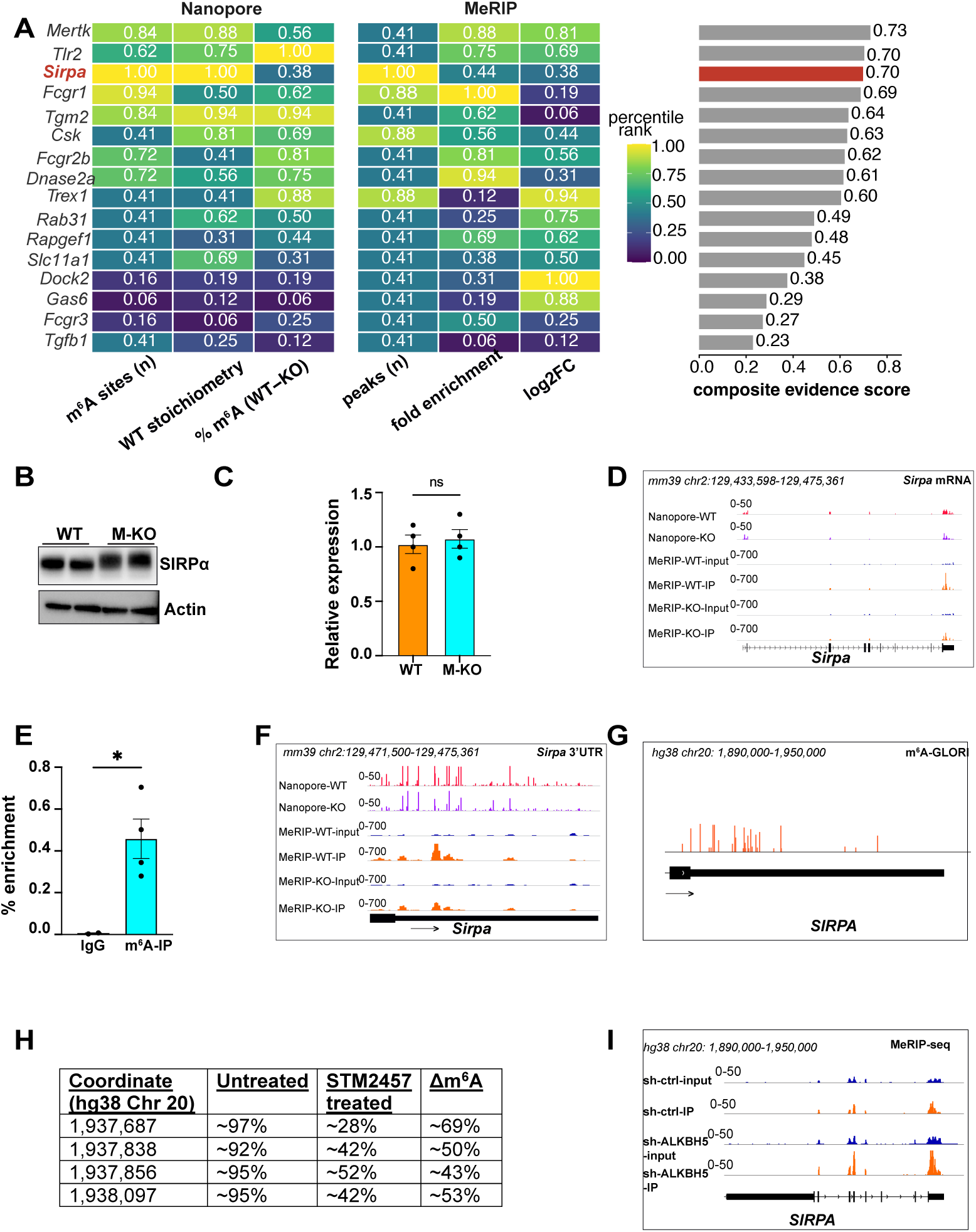
SIRPα is an m^6^A target gene. (A) Schematic of integration of MeRIP and Oxford Nanopore direct RNA sequencing data to determine m^6^A targets involved in efferocytosis and phagocytosis (B) Western blot for SIRPα in WT and M-KO BMDMs. (C) RT-qPCR for *Sirpa* in WT and M-KO BMDMs (n=2 mice per group, 2 technical replicates per mouse). (D) MeRIP-RT-PCR for *Sirpa* in BMDMs (n=2 biological replicates for IgG and 4 biological replicates for m^6^A-IP). (G) m^6^A- GLORI peak distribution on *SIRPA* in HEK293T cells. (H) Table showing the change in m^6^A at indicated positions on *SIRPA* mRNA in HEK293T cells after treatment with STM2457. (I) MeRIP peak distribution on *SIRPA* in NOMO-1 cells after treatment with shRNA-control or shRNA-ALKBH5. All data are presented as mean ± SEM. Statistical analysis in (C) and (D) was performed using unpaired t-test. *P<0.05. ns- not significant.

We next turned our attention to SIRPα, the inhibitory “don’t eat me” receptor that engages CD47 on apoptotic cells to suppress phagocytosis and efferocytosis. *Sirpa* carried the most extensive Nanopore evidence of any gene in the set, ranking first for both number of m^6^A sites and stoichiometry, and was supported by the largest number of differential MeRIP peaks. Western blot analysis showed an increase in SIRPα protein in M-KO macrophages (Fig 5B). Interestingly, this change in SIRPα protein was independent of mRNA levels, as assessed by RT-qPCR (Fig 5C).

*Sirpa* mRNA exhibited 21 Nanopore detected m^6^A sites which largely resided within 6 FDR significant MeRIP peaks, with the remaining sites within MeRIP peaks that did not pass the threshold for statistical significance. These sites were highly concentrated in the 3’UTR(Fig 5F,G), which was corroborated by MeRIP data. We validated this m^6^A modification experimentally by MeRIP-RT-PCR using poly A+ RNA isolated from murine BMDMs(Fig 5E). We also leveraged another recently developed method to detect m^6^A at single base resolution with stoichiometric information: glyoxal and nitrite-mediated deamination of unmethylated adenosines (m^6^A- GLORI)^31^. Interrogating m^6^A GLORI data from human cells, we were able to identify 18 discrete m^6^A methylated adenosine bases in the *SIRPA* 3’UTR, several of which clustered into high-density “hotspot” regions, which are considered more likely to influence transcript fate (Fig 5G). Importantly, four of these sites showed 43-69% reductions in m^6^A stoichiometry following treatment with STM2457, a selective catalytic inhibitor of the METTL3-METTL14 methyltransferase complex(Fig 5H)^19^,providing evidence that m^6^A methylation of SIRPα is conserved in mouse and human cells.

To identify the specific demethylase responsible for maintaining m^6^A at the *Sirpa* 3’UTR, we interrogated publicly available datasets profiling the interactomes of m^6^A demethylases^46^. We identified that ALKBH5 bound the 3’UTR of *SIRPA*, Critically, shRNA mediated knockdown of *ALKBH5* increased the m^6^A occupancy at this site, establishing ALKBH5 as the responsible demethylase, and suggesting that the m^6^A status of SIRPα is dynamically regulated by the opposing activities of METTL14 and ALKBH5 (Fig 5I).

We additionally identified *Gas6*, a ligand for efferocytosis receptors, as an m^6^A target gene. Nanopore and MeRIP sequencing data identified a Mettl14-dependent m^6^A site in the *Gas6* 3’UTR (Fig S5C), and *Gas6* mRNA (Fig S5D) and protein levels (Fig S5E) were significantly downregulated in *Mettl14* KO BMDMs.

Taken together, these data suggest that m^6^A directly regulates multiple elements of the efferocytosis program, including SIRPα.

## Discussion

Macrophages perform a multitude of functions in atherosclerotic plaques encompassing cholesterol handling, inflammation and tissue repair, which is tightly regulated to modulate plaque progression and regression^1^. In this study, we identify a novel epitranscriptomic regulatory mechanism controlling macrophage efferocytosis atherosclerotic plaque progression and provide genetic evidence linking m^6^A with CAD traits in humans.

Nearly 250 distinct loci have been associated with CAD risk through GWAS^34^, yet most reside in non-coding regions, making causal gene identification challenging. Molecular-QTLs (mol-QTLs) have provided a valuable framework to translate non-coding GWAS signals into causal mechanisms. Our work extends this framework to m^6^A modification, demonstrating significant enrichment of m^6^A-QTLs among CAD-associated variants and high-confidence colocalization at eight loci spanning distinct arms of atherosclerosis biology. m^6^A-QTLs could thus serve as a valuable tool for causal gene prioritization at CAD GWAS loci, complementing existing eQTL and sQTL approaches by capturing post-transcriptional regulatory mechanisms not reflected in gene expression or splicing alone.

Genome-wide m^6^A profiling has transformed our understanding epitranscriptomic regulation, yet antibody-based methods such as MeRIP seq suffer from well-recognized limitations including antibody cross reactivity with other RNA modifications, lack of single base resolution and stoichiometric information as well as variable immunoprecipitation efficiency that complicates differential quantification. Here, we leveraged Oxford Nanopore direct RNA sequencing to identify m^6^A modified bases in native mRNA, with per-site stoichiometry, and integrated these calls with MeRIP-seq to define a high-confidence set of *Mettl14* dependent m^6^A targets. This quantification of degree of modification and precise position is crucial in identification of sites with regulatory consequences and for precision approaches that write or erase m^6^A at defined positions.

Our study offers important technical information on the relative contributions of complementary m^6^A detection methods, clarifying how direct nanopore basecalling and antibody-based enrichment each shape the epitranscriptomic landscape that can be resolved. Recent advances in the Oxford Nanopore RNA004 chemistry allow direct m^6^A basecalling via a trained model (Dorado), without a matched unmodified control. This affords greater sensitivity than prior approaches using JACUSA2 basecalling in cardiomyocytes^32^, though Dorado may increase false-positive calls at low stoichiometry. This tradeoff can be mitigated by integration with MeRIP seq as we have performed.

Crucially, we were able to identify m^6^A modification of *Sirpa* mRNA in the 3’UTR using multiple orthogonal datasets including MeRIP-seq and m^6^A -GLORI and identified conservation in humans. SIRPα is the cognate receptor for CD47, an antiphagocytic “don’t eat me” ligand upregulated in tumors and atherosclerotic plaques. Inhibition of CD47 reduced atherosclerotic plaque progression in mice^14^ and vascular inflammation in humans^17^, while myeloid specific deletion of SIRPα attenuated murine atherosclerosis progression^48^, nominating the CD47-SIRPα axis as a therapeutic target in CAD. CD47-based therapeutics have been hampered by adverse events^49,50^. Attention has consequently shifted towards SIRPα itself given its more restricted expression profile on phagocytes, with multiple early-stage trials of anti SIRPα monoclonal antibodies^51^ and SIRPα fusion proteins^52^ in progress. Our identification of ALKBH5 as the putative m^6^A demethylase of SIRPα raises the intriguing concept of targeting ALKBH5 to enhance efferocytosis or phagocytosis. Indeed, recent work has also demonstrated that myeloid specific deletion of ALKBH5 reduced atherosclerosis progression and plaque inflammation^53^.

Our findings also advance understanding of the molecular regulation of efferocytosis. Physiologically, efferocytosis clears an estimated 200-300 billion apoptotic cells per day across numerous tissues^54^. Pathologically, impaired efferocytosis is causally implicated in atherosclerosis and autoimmune disease, and therapeutic enhancement of efferocytosis to reduce CAD progression is an active area of investigation^2,14,17,55^. Despite this importance, the upstream regulatory mechanisms governing efferocytosis remain poorly defined. The m^6^A -SIRPα axis we describe represents a novel homeostatic regulatory node with both physiological and pathological relevance. Finally, the identification of this axis crystalizes an important concept about the contributions of m^6^A itself in immune cells. It is thought that m^6^A may have evolved as surveillance mechanism to ‘tag’ viral or unwanted RNA for degradation^56^. Congruent with this function, enhanced efferocytosis by m^6^A could represent an adaptative mechanism to boost survival under stressful conditions.

The role of m^6^A in innate immune responses is complex, exhibiting both context and target specificity. While loss of *Mettl3* led to an increased interferon response suppressing viral propagation^57^, loss of *Mettl14* led to TLR4-driven hyperinflammation which was detrimental in the context of bacterial sepsis^21^. Interestingly, loss of *Mettl3* has been shown to dampen inflammatory responses and reduce disease susceptibility in mouse models of bacterial sepsis, colitis and gout^58,59^. Accordingly, myeloid specific deletion of *Mettl3* was also shown to attenuate inflammation and reduce atherosclerosis progression through suppression of BRAF-MAP kinase signaling^60^. This apparent dichotomy in the functions of *Mettl3* and *Mettl14* in inflammatory responses and inflammatory disease progression could be explained due to multiple different reasons. Firstly, *Mettl3* and *Mettl14* may have non-redundant regulatory targets, with different phenotypic consequences depending on stoichiometric abundance of transcripts and cellular contexts. Secondly, non-canonical functions of *Mettl3* and *Mettl14* may explain differential transcript regulation. *Mettl3* has been shown to deposit m^6^A on specialized regulatory elements such as chromosome associated regulatory RNAs (carRNAs) which can regulate transcription and chromatin state^61^. In addition, *Mettl3* and *Mettl14* have both been shown to bind chromatin in distinct patterns- *Mettl3* was shown to be associated with loci marked by the repressive histone mark H3K9Me3^62^, while *Mettl14* associated with both the repressive histone mark H3K27Me3 and the active histone marks H3K27Ac and H3K4Me3^63^. These differential chromatin binding patterns of the writer proteins could also explain divergent phenotypes in their absence.

Our study is not without limitations. A key limitation of our genetic analysis is the ancestry mismatch between the m^6^A-QTLs, which were mapped in individuals of Yoruba (YRI) African ancestry, and the CAD GWAS, which was conducted in individuals predominantly of European ancestry. Differences in linkage disequilibrium (LD) structure between these populations may result in more conservative estimates of colocalization due to an inflation in H3 (evidence of distinct variants) rather than H4 (shared variants). It is possible that the increased atherosclerotic plaques in *Mettl14* M-KO mice are also due to additional mechanisms other than impaired efferocytosis. We also cannot exclude m^6^A independent effects of *Mettl14* including changes in caRNAs and chromatin accessibility.

## Data Availability

Public data used was obtained from NCBI GEO Accession numbers: GSE125377 (MeRIP seq from 60 YRI samples for m^6^A-QTL analysis) GSE153511 (MeRIP seq from murine BMDMs), GSE210563 (m^6^A GLORI in HEK293T cells). GWAS summary statistics for CAD were obtained from (http://www.cardiogramplusc4d.org/), summary statistics for m^6^A- QTL analysis were obtained from https://doi.org/10.5281/zenodo.3870952. Oxford Nanopore direct RNA sequencing will be uploaded on NCBI GEO.

## Acknowledgements

A.K. is supported by an American Heart Association Predoctoral Fellowship (25PRE 1359710), X.L. is supported by an American Heart Association postdoctoral fellowship (26POST 1564949), E.R.S. was supported by an American Heart Association Postdoctoral Fellowship (24POST 1183446), Z.Z. is supported by an American Heart Association Career Development Award (25CDA 1453023), D.W. is supported by the National Institutes of Health K99 HL175030-01A1. Y.A.C. is supported by the National Institutes of Health R01 HL185164 and RC2 DK144739 , W.L. is supported by the National Institutes of Health R01 CA290720, CA193466, RC2 DK144389, HL180397. X.H. is supported by the National Institutes of Health R01 AI175554, U19 AI162310, R01 HL188168, R01 AG095819, C.H. is supported by the National Institutes of Health R01 HG006827, R01 HL155909, R01 ES038076, RC2 DK139552, R33 CA309701 and the Howard Hughes Medical Institute, T.S. is supported by the National Institutes of Health R01 HL139549, R01 HL149766, R01 HL180397, R01 HL185164 and RC2 DK144739. The authors would also like to acknowledge the UCLA Immune Assessment Core, UCLA Flow Cytometry Core (supported by National Institutes of Health P30 CA016042 and 5P30 AI028697, Jonsson Comprehensive Cancer Center, UCLA AIDS Institute, David Geffen School of Medicine at UCLA, UCLA Chancellor’s Office, UCLA Vice Chancellor’s Office of Research) CNSI Advanced Light Microscopy and Spectroscopy Core (RRID:SCR_022789) (Funding support from NIH Shared Instrumentation Grant S10OD02017, NSF Major Research Instrumentation grant CHE-0722519) and the UCLA Lipidomics Core.

## Methods

### m^6^A-QTL integration

First, we performed a binary annotation enrichment analysis. SNPs in the Coronary Artery Disease GWAS were annotated as m^6^A QTLs if their chromosomal position matched a significant m^6^A QTL (FDR < 10%). Enrichment was quantified using Fisher’s exact test comparing the proportion of m^6^A QTL SNPs among genome-wide significant CAD variants (p < 5×10⁻⁸) versus non-significant variants. To assess the robustness of this enrichment across significance thresholds, the analysis was repeated at five p-value cutoffs (p<5x10^-8^, 1x10^-8^, 1x10^-6^, 1x10^-5^, 1x10^-4^, 1x10^-3^). To visualize the enrichment across the full allelic spectrum, quantile-quantile (Q-Q) plots were generated stratifying CAD GWAS p-values by m^6^A QTL status, with 100,000 non- m^6^A QTL SNPs randomly sampled for computational efficiency.

Pre-trained LASSO regression models of m^6^A levels from genotype data were obtained from^33^ GWAS effect alleles and test statistics were harmonized to the reference and effect alleles of the LASSO models prior to z score computation.

To validate the TWAS-colocalization findings and account for LD between nearby genes, be applied the causal TWAS (cTWAS) framework. We supplied pre-computed TWAS z-scores directly and ran the no-LD version of cTWAS (ctwas_sumstats_noLD), which assumes at most one causal signal per LD region. Model parameters (prior inclusion probabilities and effect size variances) were estimated with expectation-maximization (EM) and converged after 7 iterations.

### Oxford Nanopore Direct RNA sequencing

Poly(A)-selected RNA was isolated from Bone Marrow Derived Macrophages isolated from WT or M-KO mice. Sequencing was performed on Oxford Nanopore RNA004 chemistry using PromethION FLOW-PRO004RNA flow cells.

### Animals and diets

All mice used in this study were on the C57BL/6 background. While our *in-vivo* studies were performed only in male mice, *in-vitro* studies were performed in primary macrophages isolated from both male and female mice. Mettl14^f/f^ mice were obtained as previously described^23^. Mice were housed in a temperature-controlled vivarium with a 12-hour light-dark cycle. Mice were fed either a standard rodent chow or Western Diet with 0.2% Cholesterol (Research Diets, D12079B). *Ldlr*^-/-^ mice were purchased from the Jackson Laboratory (JAX, Cat. No. 002207). For the atherosclerosis study, mice were fasted for 4 hours prior to sacrifice. After bone marrow transplantation, mice were housed in a room with autoclaved cages and bedding.

### Cell culture and Treatments

Jurkat cells were originally obtained from ATCC. Peritoneal macrophages were cultured in Dulbecco’s Modified Eagle’s Medium (DMEM), 10% Fetal Bovine Serum (FBS) and 1% Penicillin- Streptomycin. BMDMs were cultured in DMEM, 20% FBS, 10% CMG supernatant, 1% Penicillin- Streptomycin, 1% L-glutamine and 0.5% Sodium pyruvate. Jurkat cells were cultured in Roswell Park Memorial Institute Medium (RPMI), 10% FBS and 1% Penicillin-Streptomycin. Peritoneal macrophages were cultured in DMEM, 10% FBS and 1% Penicillin-Streptomycin.

For acLDL and oxLDL treatment, cells were sterol starved for 6-12 hours prior to the experiment in sterol depletion medium: culture medium with 2% lipoprotein deficient serum (Sigma S5519) supplemented with 5µM Simvastatin and 100µM mevalonate. After the starvation period, cells were treated with the indicated lipoproteins for the given period. Acetylated LDL and Oxidized LDL were obtained from Invitrogen (L35354, L34358 respectively)

### Bone marrow derived macrophage isolation

After sacrificing Mettl14^f/f^ and Mettl14^f/f^ LysM^Cre^ mice, the skin, muscle and connective tissue overlying the hindlimbs was dissected off. Bone marrow was flushed out of the femur and tibia using cold PBS. After washing with RBC lysis buffer, bone marrow cells were cultured in the medium mentioned above. At day 3 of culture, the medium was changed. BMDMs were used for experiments between day 6-8 of differentiation. For *in-vitro* efferocytosis experiments, the cells were cultured in heat-inactivated FBS.

### Peritoneal macrophage isolation

Mettl14^f/f^ and Mettl14^f/f^ LysM^Cre^ mice were injected with 1ml thioglycolate broth intraperitoneally. Four days later, mice were sacrificed and cells were harvested by peritoneal lavage with cold PBS.

After RBC lysis, cells were plated in the medium indicated above and experiments were performed the following day.

### Gene expression and immunoblot

For gene expression analysis, total RNA was isolated using TRIzol reagent (Invitrogen). 500ng-1 µg RNA was subjected to reverse transcriptase reaction using a homemade Reverse Transcriptase enzyme. Quantitative PCR analysis was performed using a BioRad SYBR-green system and Quant Studio PCR machine. RT-qPCR analysis was performed using the delta-delta CT method with Cyclophilin A (*Ppia*) as the housekeeping gene. For immunoblotting, samples were collected in Radioimmunoprecipitation assay (RIPA) buffer supplemented with protease and phosphatase inhibitors, following which protein concentration was determined using a BCA assay. Denaturation was performed using a Lithium dodecyl sulfate based buffer (NuPAGE-LDS) supplemented with 5% Beta-mercaptoethanol. Equal amounts of protein were loaded on a NuPAGE Bis-Tris Gel (Invitrogen NP0322), and SDS-PAGE was performed using MOPS running buffer, followed by a wet transfer. Blocking was performed using 5% non-fat milk solution in TBST at room temperature for 1 hour, primary antibody incubation was performed at 4°C overnight, and secondary antibody incubation was at room temperature for 1 hour. All loading controls were from the same membrane as the original samples, and stripping was performed using stripping buffer (Fisher, 46430) when the loading control and antigen of interest were of similar molecular weights. Immunoblotting for GAS6 was performed in non-reducing conditions.

### Atherosclerosis study and lesion analysis

Bone marrow cells were collected from Mettl14^f/f^ and Mettl14^f/f^ LysM^Cre^ mice. Approximately 5x10^6^ bone marrow cells were injected intravenously through the tail vein to *Ldlr*^-/-^ mice lethally irradiated with 900 rads. Following transplantation, mice were allowed to recover for four weeks with antibiotic (Baytril) water, after which the mice were fed with Western Diet (Research Diets, D12079B) for 18 weeks. The aorta was harvested and atherosclerosis lesions were quantified as previously described^64^. Briefly, the left ventricle was perfused via a cannula, following which the aorta was fixed by perfusion with formal sucrose. After perfusion, the aorta was exposed, adventitial tissue was trimmed. Atherosclerosis quantification of Sudan IV-stained aortae was quantified using a Zeiss Stemi 508 microscope as the percentage of entire aortic area with atherosclerotic lesions. Aortic roots were harvested in Optimal Cutting Temperature (OCT compound) and stored in -80°C. Serial sections at the level of the aortic valves were harvested and used for histological analysis. Genomic DNA was isolated from peripheral blood using the Wizard Genomic DNA Purification Kit (Promega)

### Plasma lipid and cytokine measurement

Plasma cholesterol and triglycerides were measured using Fujifilm Wako kits as per the manufacturer’s instructions. Plasma levels of TNFα, IL-6, RANTES and MCP-1 were measured using a Luminex ELISA kit at the UCLA Immune Assessment Core. Plasma levels of IL-18 were measured using ELISA (R&D Systems, cat no 7625).

### Lipidomics

WT and M-KO peritoneal macrophages were treated with 75 µg/mL acLDL for 12 hours. Lipid extraction and Lipidomics was performed at the UCLA Lipidomics core. In brief, lipid extraction was performed using a modified Bligh and Dyer extraction technique with a Lipidyzer Internal Standard Mix (13 lipid classes, AB Sciex). Organic phases were then dried and resuspended in solvent (1:1 methanol-dichloromethane, with 10mM ammonium acetate). Lipid species were quantified on the Sciex Lipidyzer with SelexION differential mobility tuning, and data were processed using the Lipidyzer software. Quantitative values were normalized to cell number.

### Atherosclerosis analysis

For Hematoxylin and Eosin staining, formalin fixed slides were stained with hematoxylin (Richard- Allan Scientific, 7211). After washing in ethanol, slides were immersed in alcoholic eosin solution, washed and then mounted. Trichrome staining was performed using a kit (Sigma HT15) according to the manufacturer’s instructions. For Oil Red O staining, formalin fixed slides were stained with fresh Oil Red O solution and counterstained with hematoxylin.

### Foam cell assay

WT and M-KO peritoneal macrophages were treated with 100 µg/mL oxidized LDL cholesterol for 72 hours. Oil Red O staining was performed as mentioned above.

### Immunofluorescence and Immunohistochemistry

Immunofluorescence staining was performed on frozen aortic root sections. Sections were fixed in 4% paraformaldehyde and permeabilized in 0.1% Triton X-100 solution. Sections were then blocked in 3% Bovine Serum Albumin (BSA) solution and then incubated with the indicated antibodies at 4° Celsius overnight. Secondary antibody incubation was performed at room temperature in the dark for 1 hour. Slides were then stained with Hoechst 33342 and finally mounted using ProLong Gold Antifade mounting reagent. Immunohistochemistry staining on frozen aortic root sections was performed using the VECTASTAIN ABC-AP kit as per the manufacturer’s instructions. TUNEL staining was performed on frozen aortic root sections using the Click-iT Plus TUNEL assay kit (Invitrogen, C10617) as per the manufacturer’s instructions.

### Dexamethasone-Thymus study

WT and M-KO mice were injected with 250 µg water soluble Dexamethasone (Sigma, D2915) in PBS. Eighteen hours later, the thymi were harvested by careful dissection under a light microscope and were weighed. Cellular suspensions were obtained by dissociation and cell counts were obtained using automated cell counting (Invitrogen Countess). Cells were then stained with Annexin-V and flow cytometry was performed using Attune II (Thermo Fisher).

### In-vitro efferocytosis

Jurkat cells were labeled with green CMFDA (MedChemExpress) as per the manufacturer’s instructions. Apoptosis was induced in Jurkat cells using an 8W UV lamp (256nm, AnalytikJena) for 10 minutes, followed by culture for 3 hours. Labeled apoptotic Jurkat cells were co-incubated with WT and M-KO BMDMs in a 1:5 (BMDM to apoptotic cell) ratio for 1 hour. After one hour, the cells were carefully washed with PBS to remove unbound apoptotic cells. The cells were fixed in 4% paraformaldehyde and stained with Wheat Germ Agglutinin conjugated to Texas Red.

### Imaging and Quantification

Hematoxylin and Eosin staining, Oil Red O staining, Trichrome staining, CD68 immunohistochemistry and foam cell assay were imaged using a Zeiss Vert A1 microscope. Image quantification was performed using ImageJ.

Immunofluorescence and *in-vitro* efferocytosis images were imaged using a Leica SP8 confocal microscope at the UCLA Advanced Light Microscopy and Spectroscopy Core facility. Image acquisition was performed using Z-stack imaging and image processing was performed using maximal projection. Image analysis was performed using LasX software (Leica).

### MeRIP-PCR

mRNA was isolated from BMDMs using Dynabeads mRNA Direct purification kit (Invitrogen). 2 µg of mRNA per replicate was chemically fragmented using NEBNext Magnesium RNA Fragmentation Module (New England Biolabs, E6150) for exactly 4 minutes at 94 degrees Celsius. M6a-IP was performed using the Magna MeRIP kit (Sigma 17-10499). Eluted mRNA was reverse transcribed using SuperScript III reverse transcriptase (Invitrogen) and qPCR was performed as previously described.

---

**Figure S1:**
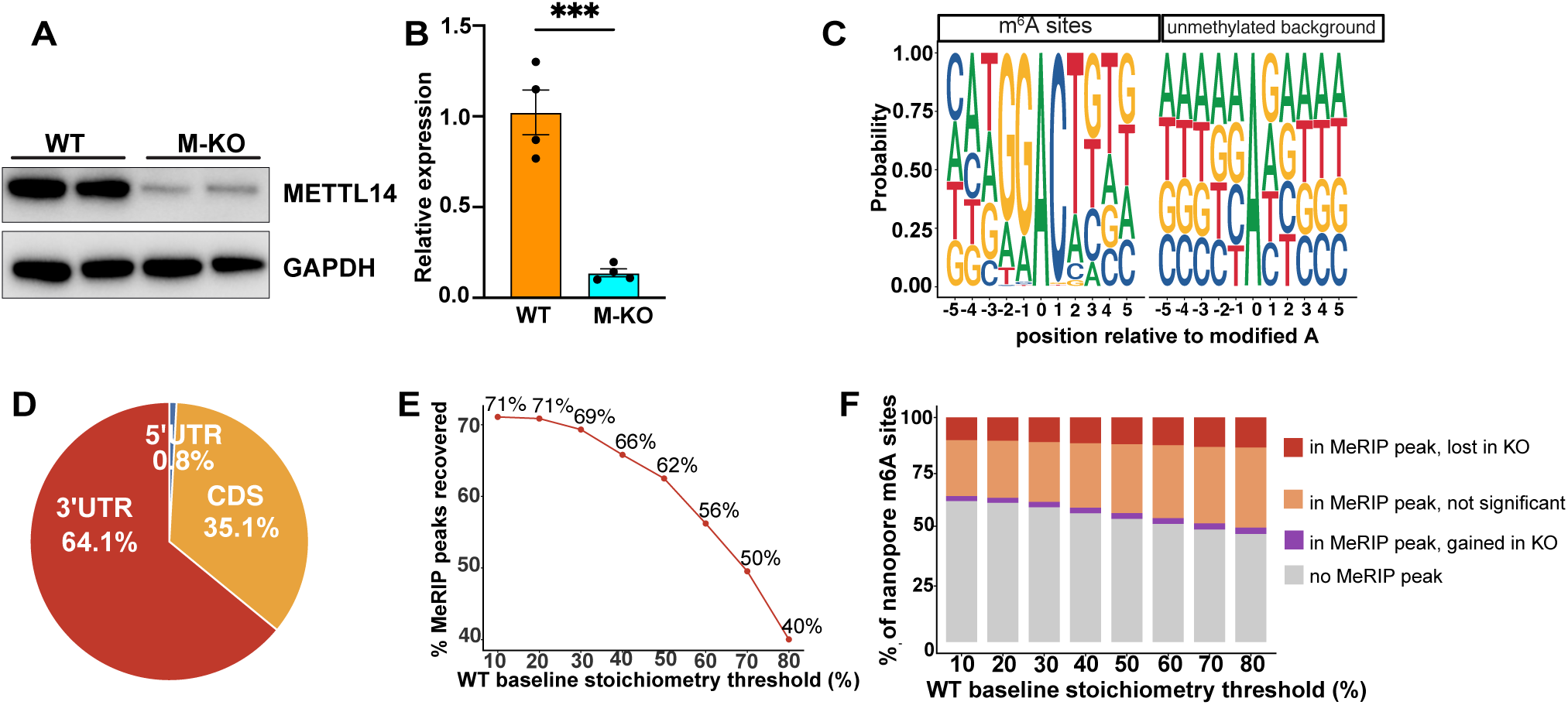
Oxford Nanopore direct RNA sequencing nominates m^6^A target genes. (A) Western blot for METTL14 in WT and M-KO bone marrow derived macrophages (n=2 mice per group). (B) qPCR for *Mettl14* in WT and M-KO BMDMs (n=2 mice per group, 2 technical replicates per sample). (C) Motif enrichment at putative m^6^A motifs in Nanopore sequencing data. (D) Distribution of Nanopore m^6^A sites. (E) Percentage of MeRIP peaks identifed by Nanopore sequencing data at various WT baseline stoichiometric thresholds. (F) Percentage of Nanopore m6A sites found in MeRIP sequencing data. Statistical signficance in (B) was assessed using student’s unpaired t-test. Data in (B) represent mean ± SEM. Data in (F) represent FDR<0.05. ***p<0.00. UTR= untranslated region, CDS= coding DNA sequence, MeRIP= m^6^A immunoprecipitation and sequencing.

**Figure S2:**
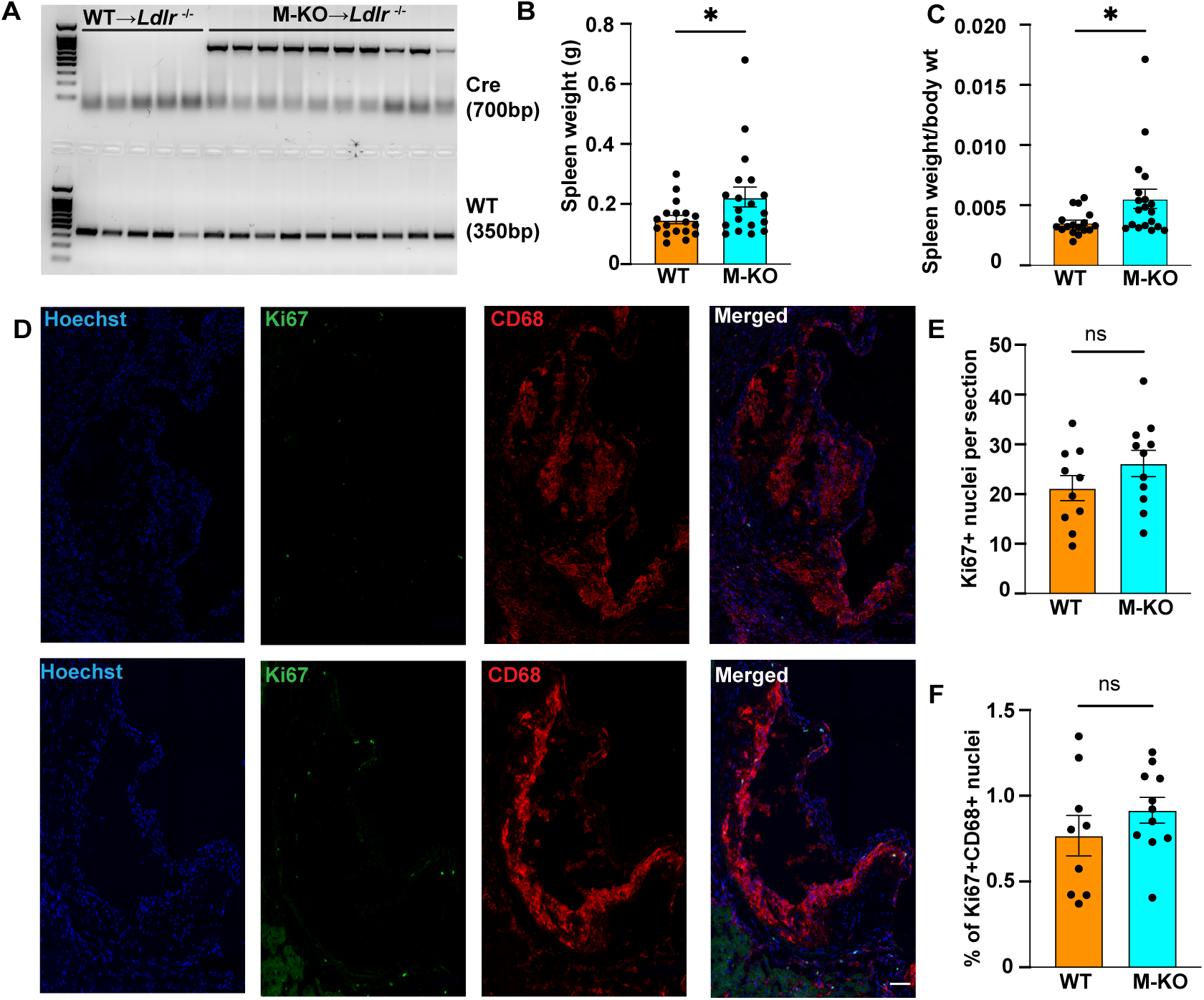
Impact of loss of myeloid *Mettl14* on atherosclerotic plaque characteristics . (A) PCR for Cre or WT alleles isolated from genomic DNA isolated from peripheral blood of *Ldlr*^-/-^ mice transplanted with WT or M-KO bone marrow (n=5 WT, 10 M-KO). (B) Spleen weight of BMT mice (n=18 WT, 19 M-KO). (C) Spleen weight of BMT mice normalized to body weight (n=17 WT, 19 M-KO, one sample from WT formally excluded as an outlier using Grubb’s test P<0.05). (D) Immunofluorescence of aortic roots from BMT mice for Ki67 (green) and CD68 (red), scale bars represents 100μm . (E) Quantification of number of Ki67 nuclei per section (n=10 WT, 11M-KO). (F) Quantification of percentage of Ki67 CD68+ nuclei (n=9 WT, 11 M-KO).All data are presented as mean ± SEM. Statistical analyses in (B), (C), (E), (F) were performed using student’s t-test. *P<0.05. ns- not significant.

**Figure S3:**
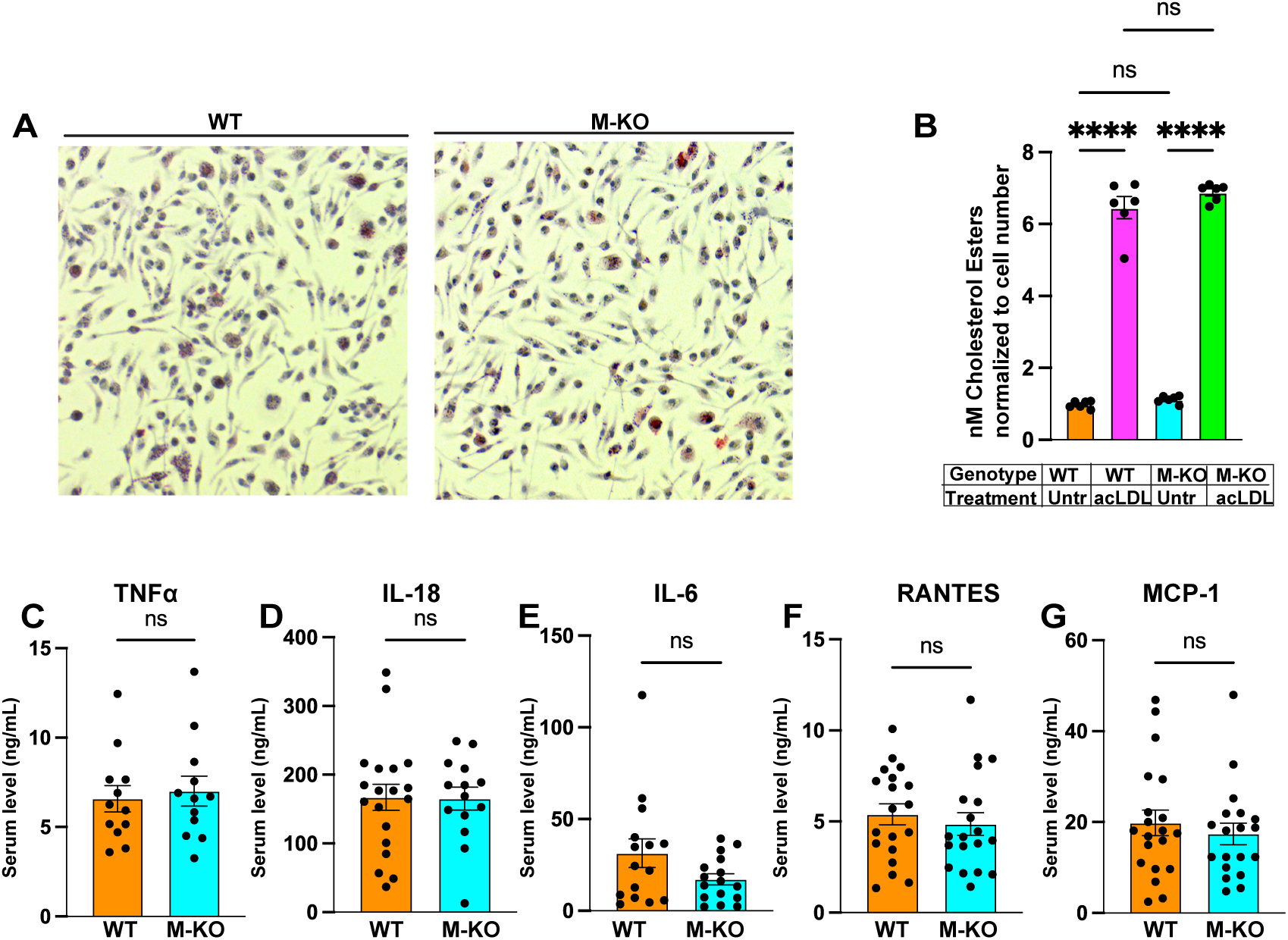
Loss of myeloid *Mettl14* does not alter foam cell formation or systemic inflammation in atherosclerosis. (a) Representative image of Oil-Red-O staining in WT and M-KO peritoneal macrophages treated with 100µg/mL oxLDL for 72hr. (b) Cholesterol ester quantification in WT and M-KO peritoneal macrophages treated with 75µg/mL acLDL for 12hr. (c) TNFα levels measured using ELISA in the serum of BMT mice. (d) IL-18 levels measured using ELISA in the serum of BMT mice. (F) RANTES levels measured using ELISA in the serum of BMT mice. Statistical significance in (b) was assessed using one-way ANOVA with Tukey’s multiple comparisons test.Statistical significance in (c), (d), (e), (f), and (g) was assessed using unpaired student’s t test. All data are presented as mean ± SEM. ****p<0.0001. ns- not significant. RANTES- Regulated upon Activation, Normal T cell Expressed and Secreted. MCP-1= Monocyte Chemoattractant Protein-1. TNFα= Tumor Necrosis Factor Alpha.

**Figure S4:**
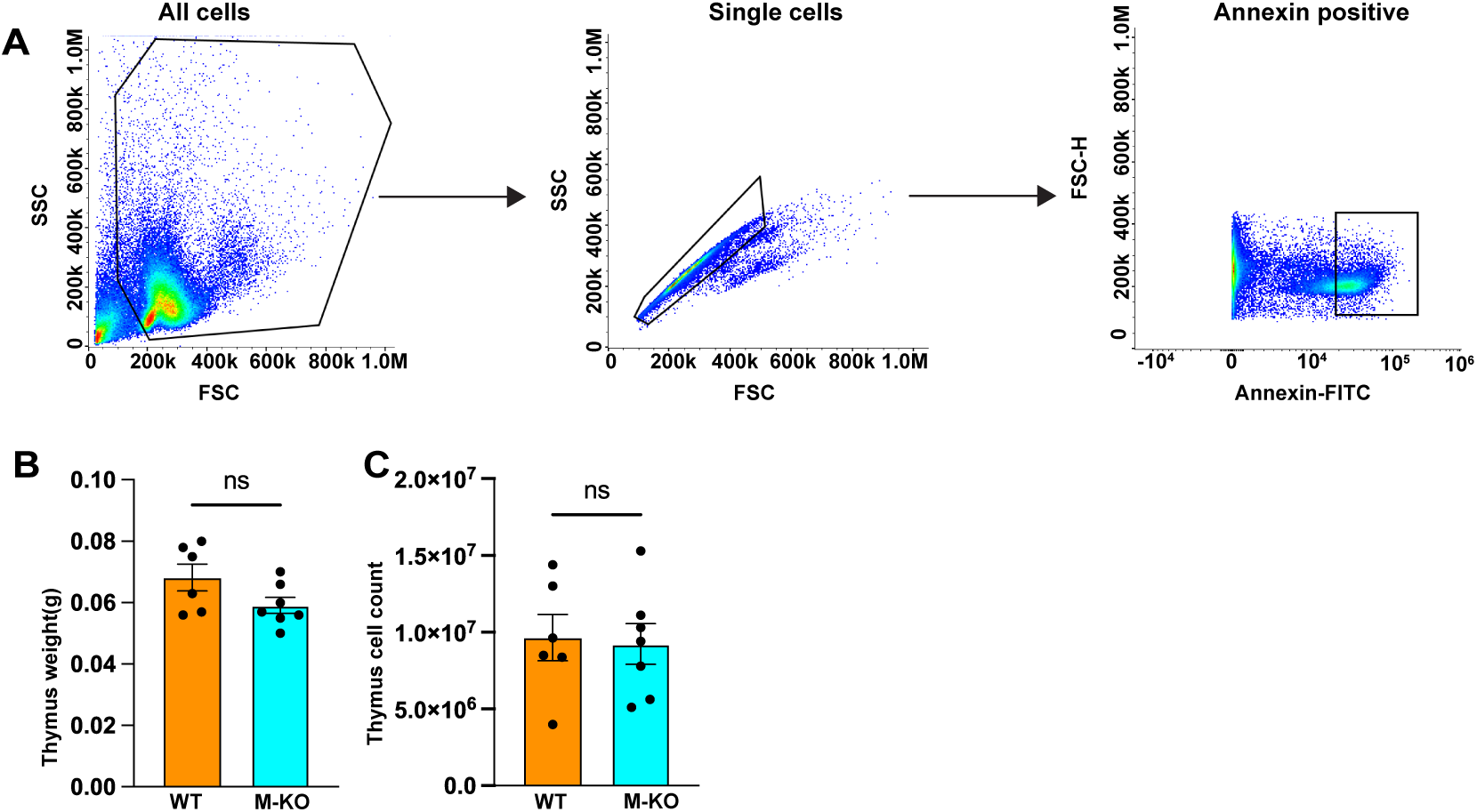
Loss of myeloid Mettl14 leads to impaired efferocytosis. (A) Gating strategy for flow cytometry. (B) Thymus weight in WT and M-KO mice injected with PBS (n=6 WT, 7 M-KO). (C) Thymus cell count in WT and M-KO mice injected with PBS (n=6 WT,7 M-KO). All data are presented as mean ± SEM. Statistical analysis in (B) and (C) was performed using unpaired t-test.

**Figure S5:**
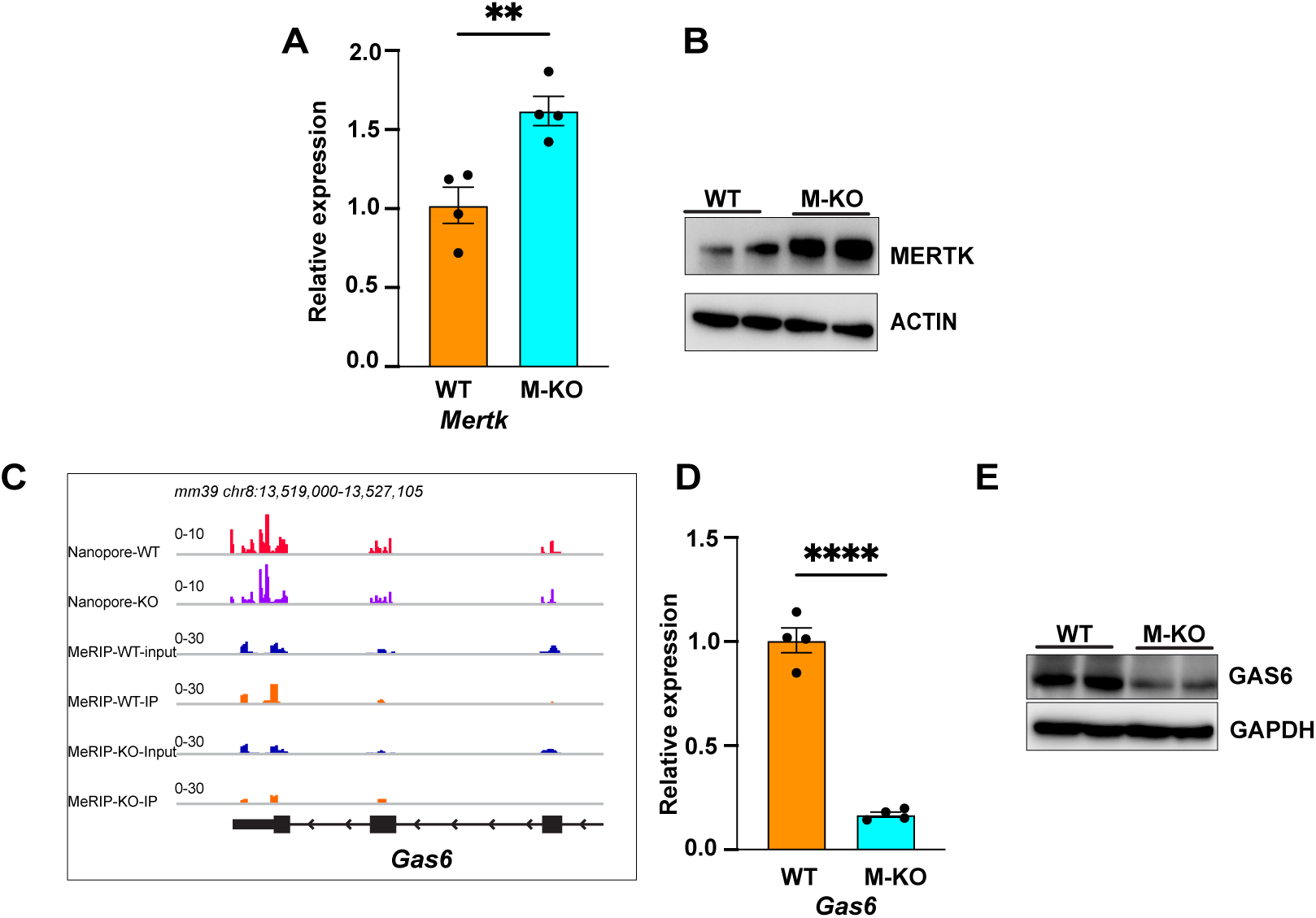
m^6^A targets include transcripts involved in efferocytosis. (A) RT-qPCR for *Mertk* in WT and M-KO BMDMs (n=2 mice per group, 2 technical replicates per mouse). (B) Western blot for MERTK in WT and M-KO BMDMs (n=2 mice per group). (C) Nanopore and MeRIP seq data showing the m^6^A peaks on the *Gas6* transcript. (D) RT-qPCR for *Gas6* in WT and M-KO BMDMs (n=2 mice per group, 2 technical replicates per mouse). (E) Western blot for Gas6 protein in WT and M-KO BMDMs (n=2 mice per group). Data are presented as mean ± SEM. P values were determined by unpaired-t test, **P<0.01, ****P<0.0001

